# Regional and fine-scale local adaptation in salinity tolerance in *Daphnia* inhabiting contrasting clusters of inland saline waters

**DOI:** 10.1101/2023.08.11.552416

**Authors:** Kristien I. Brans, Csaba F. Vad, Zsófia Horváth, Luca Santy, Kiani Cuypers, Robert Ptacnik, Luc De Meester

**Author notes:** Corresponding author: Csaba F. Vad. These authors contributed equally to this study (Kristien I. Brans, Csaba F. Vad, Zsófia Horváth).

## Abstract

Understanding the spatial scales at which organisms can adapt to strong natural and human-induced environmental gradients is important. Salinisation is a key threat to biodiversity, ecosystem functioning, and the provision of ecosystem services of freshwater systems. Clusters of naturally saline habitats represent ideal test cases to study the extent and scale of local adaptation to salinisation. We studied local adaptation of the water flea *Daphnia magna*, a key component of pond food webs, to salinity in two contrasting landscapes - a dense cluster of sodic bomb crater ponds and a larger-scale cluster of soda pans. We show regional differentiation in salinity tolerance reflecting the higher salinity levels of soda pans versus bomb crater ponds. We found local adaptation to differences in salinity levels at the scale of tens of metres among bomb crater pond populations but not among geographically more distant soda pan populations. More saline bomb crater ponds showed an upward shift of the minimum salt tolerance observed across clones and a consequent gradual loss of less tolerant clones in a nested pattern. Our results show evolutionary adaptation to salinity gradients at different spatial scales and fine-tuned local adaptation in neighbouring habitat patches in a natural landscape.

## INTRODUCTION

In heterogeneous natural landscapes, organisms often face steep environmental gradients, while human-induced environmental changes also often result in steep gradients of stressors, such as in the case of urbanisation or intensive agriculture [1–3]. There is evidence that evolutionary trait change and adaptation can contribute to buffering ecological gradients and change in terms of population and community features [4–6]. In this context, the spatial scale at which evolutionary trait differentiation and local adaptation occurs is an important question [7], both in terms of the capacity of populations to survive but also in terms of resilience of community composition, diversity and ecosystem functioning. Theory predicts that the occurrence of local adaptation depends on the selection-migration balance [8–11]. It can therefore be expected that the likelihood of local adaptation is higher when selection gradients are stronger and gene flow is not so high as to homogenise genetic variation across habitats. The latter suggests that local adaptation may depend on geographic distances. Yet, there are several mechanisms that interfere with this general inference and allow patterns of local adaptation at very small spatial scales [7].

Salinity is a key abiotic gradient in aquatic systems. Inland waters cover a wide range of salinity levels [12], with a strong structuring role on biodiversity, community composition, and ecosystem functioning [13–15]. Naturally saline waters are globally widespread, especially in dry climatic regions [12,16] and in areas dominated by mineral-rich soils, such as soda pans in Central Europe [17]. These habitats have a high conservation importance, by hosting unique salt-adapted communities and providing feeding grounds for waterbirds [16,18,19]. Recently, there has been an increased focus on saline lakes in the context of anthropogenic freshwater salinisation, a process driven by mining effluents, road deicing, unmanaged irrigation and climate change, and emerging as a global threat to freshwater resources [20–22]. Understanding to what extent species can adapt to increasing salinities and how this relates to landscape features such as geographic distances among patches and the steepness of the salinity gradients thus becomes increasingly important in our efforts to predict responses to global climate change.

Salinisation leads to strong changes in community composition through species sorting. Communities of inland saline waters often are a nested subset of freshwater communities [14,23,24], whereas species replacement becomes more prominent at higher salinities [13,24,25]. Parallel to environmental sorting structuring metacommunities, salinity changes can also exert strong selection pressures impacting the evolution of local populations and consequently metapopulation structure [26,27]. There is evidence for local adaptation to salinity gradients in multiple taxonomic groups including fish, amphibians, and insects [28–30]. For example, populations of the amphibian *Hyla cinerea* in anthropogenically salinising coastal wetlands evolved an increased salinity tolerance compared to freshwater populations, showing a higher fecundity and increased hatching success and tadpole survival upon salt-exposure [29].

Clusters of naturally saline aquatic habitats provide excellent model systems to explore how, to what extent and at which spatial scale freshwater species can adapt to salinity gradients. The sodic waters in the Pannonian Biogeographic Region are inland alkaline saline waters, with an ionic composition typically dominated by sodium (Na^+^) and carbonates (CO_3_^2–^and HCO_3_^−^ [31]). They are naturally saline waters in this region, where their occurrence is related to climatic and geological drivers [31,32]. Their salinity strongly varies in space and time [14,33]. Soda pans dry out annually and thus exhibit a wide range in salinities, from sometimes freshwater in early spring to high values and, in extreme cases, hypersalinity (> 50 g L^−1^) at the end of the growing season [17,19]. Other sodic habitats in the region, such as bomb crater ponds, are less extreme in their salinity, with not all ponds drying up every year [34].

*Daphnia magna* is a freshwater zooplankton species that typically occurs in small ponds and shallow lakes. Because of its relatively broad salinity tolerance compared to many other cladoceran zooplankton species [14,35,36], it thrives in sodic waters, also because they are generally fishless and turbid, which reduces the vulnerability of this large-bodied cladoceran to predation [34,37]. The widespread occurrence and high abundances of *D. magna* in sodic waters [14] and the fact that these habitats can widely differ in salinity offers a unique model system to study the degree of local adaptation and to reveal at which spatial scale it occurs. *Daphnia* is a key component of many inland water food webs by linking primary production to higher trophic levels [38,39], and has been shown experimentally to have the capacity to adapt to salinisation [40,41]. One of the best studied examples involves obligate parthenogenetic lineages of *D. pulex*, shown to be adapted to salinity gradients by strong clonal sorting, with clear differences in salinity tolerance between clones originating from low- and high-salinity habitats [42,43].

Here, using a quantitative genetics common garden approach, we test for the salinity tolerance of 126 clones from 21 *D. magna* populations occurring in the same biogeographic region. We use populations from both soda pans (Seewinkel region, Austria) and sodic bomb crater ponds (Kiskunság region, Hungary) as model systems to quantify the degree and spatial scale of local adaptation to salinity (Figure 1). Given the higher salinity levels of soda pans compared to bomb crater ponds, we expect regional differences, with the *D. magna* populations isolated from the soda pans showing a higher salinity tolerance. With respect to local adaptation, these two landscapes provide a highly interesting contrast. Geographic distances among soda pans (on average 7000 m) are much higher than among the bomb crater ponds (on average 300 m), and thus one might expect more opportunities for local adaptation in soda pans than in bomb crater ponds. Yet, other local and landscape features may lead to the opposite prediction. As soda pans show strong intra-year variation in salinity and as dispersal can be facilitated by both waterbirds and strong winds among them [44,45], one may expect no local adaptation in these habitats. In contrast, even though bomb crater ponds are much closer to each other, we might expect local adaptation, because they systematically differ in salinity, are more sheltered from wind, and not intensively frequented by birds. We also aimed to explore how local habitat salinities determine clonal variation in salinity tolerance of populations in the two study systems. We test the hypothesis that, similar to patterns seen in species occurrences along salinity gradients, salinity tolerances of the populations from more saline habitats are subsets of the salinity tolerances of populations from habitats with lower salinities. This pattern would reflect that genotypes characterised by lower salinity tolerances are weeded out in the high salinity habitats.

**Figure 1.**
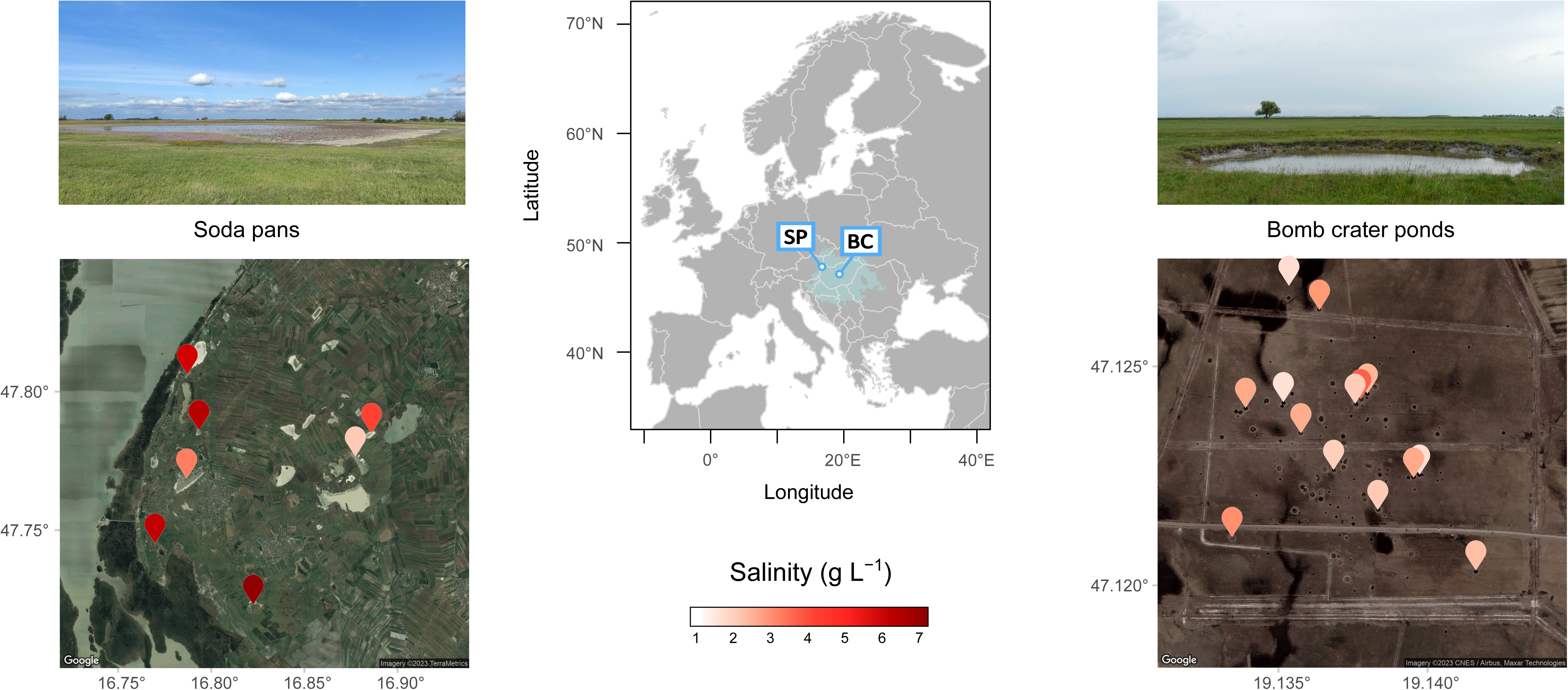
The locations of the two study systems (SP: soda pans, BC: bomb crater ponds) in the Pannonian Biogeographic Region (light blue) in Europe. Colour coding visualises mean salinity values across years and seasons of the seven studied soda pans in Austria and fourteen bomb crater ponds in Hungary. Detailed information on local habitat characteristics is given in Supplementary Material, Table S1.

## MATERIAL AND METHODS

### Study systems

Soda pans are shallow, saline, and fishless water bodies, which generally dry out in summer. Within Europe, their occurrence is restricted to the Pannonian Plain; the lowlands of Hungary, eastern Austria, and northern Serbia. The predominant environmental gradient, salinity, is a strong selection factor, and only a limited set of highly tolerant species account for the largest part of zooplankton biomass [14]. This includes *D. magna*, which is the most widespread species in these habitats [37]. The Seewinkel region in the northwestern part of the Pannonian Plain is part of a UNESCO World Heritage Site. It hosts a cluster of 34 soda pans, formed in a confined region of approximately 200 km^2^ (Figure 1; [44]). Their salinity represents a wide gradient ranging from 1.4 to 13.0 g L^−1^ [17]. They are spatially discrete ecosystems without natural in- or outflow; the nearest distance to another pan ranges from 100 m to 18 000 m with an average distance of 7000 m among habitats [17]. The surface area of the pans varies from 2000 to 973 000 m^2^. The pans have a high surface to depth ratio and are very exposed to wind. The region experiences very constant unidirectional winds, which result in a directional spatial imprint on zooplankton metacommunity structure [44].

In the Kiskunság region of Central Hungary, there is a dense cluster of more than 100 bomb crater ponds situated within an area of ∼1 km^2^, created by mistargeted bombing during World War II (Figure 1). They hold sodic water and exhibit a gradient in salinity between 1.0–5.6 g L^−1^ [34]. Their surface area varies between 7 and 113 m^2^. The average distance (calculated based on pond centre points) between them is 300 m and ranges from 9 to 860 m, which in some cases results in ponds that are only a mere 2-3 m apart (effective distance). *D. magna* is one of the dominant species in these ponds, being present in 66.7% of the ponds across a wide salinity gradient [34].

### Sampling and culturing of *D. magna*

During the period 10-13 April 2018, a total of 21 *D. magna* populations were sampled; 14 bomb crater pond and seven soda pan populations (Figure 1, Supplementary Material, Table S1). The distances among these subsets of ponds ranged from 17 to 842 m (average distance: 333 m) in case of the bomb crater ponds, and 1201 to 9869 m (average: 6217 m) for soda pans. During sampling, we also measured conductivity and other environmental variables (pH, water depth, total suspended solids and chlorophyll *a* concentrations) following the methods described in [14] and [34] (Supplementary Material, Table S1). We converted conductivity to salinity by applying a multiplying factor of 0.774 established for soda waters [14,46]. At the time of sampling, pond salinities ranged from 0.69 to 3.56 g L^−1^ in bomb crater ponds and from 1.36 to 7.07 g L^−1^ in soda pans (Supplementary Material, Table S1). As a more general measure of salinity, we collected available data from spring and summer salinity values reported from the same habitats ([14,34], and own unpublished data, listed also in Supplementary Material, Table S1). These data were first averaged per season and pond, and then for each pond, we also calculated a general mean (the mean of mean spring and mean summer values). We used these mean values per pond to illustrate general differences in salinities irrespective of seasons (Figure 1).

Between 8 and 20 *D. magna* individuals were isolated from each population and cultured as clonal lineages. Of these, six clonal lineages from each population were randomly picked out to be used in the common garden experiments, resulting in 21 populations x 6 lineages = 126 lineages. All clonal lineages were expected to be unique clones based on a genetic analysis using 13 microsatellite markers carried out on multiple soda pan and bomb crater populations that were sampled in April 2014, where we observed that all individuals differed in multi-locus genotype ([47]; Horváth Z. pers. comm.). Based on this, we expect that there were none or only very few genotypes that were represented by two or more individuals. Parthenogenetically produced offspring of each clone were used in experiments after purging maternal effects over at least three generations (details are provided in the Supplementary Material, section Material and Methods).

### Salinity tolerance experiments

In an effort to cover the natural range of salinities in the studied landscapes, we established 8 different salinity treatments containing 0, 3, 4, 5, 6, 7, 8, 9 g L^−1^ NaHCO_3_. We used NaHCO_3_ dissolved in dechlorinated tap water, as it is the most frequent type of salt determining the salinity of sodic waters of the Pannonian Plain [17]. Salt solutions were prepared daily for use in the experiment. Conductivity as a proxy indicated that salinity consistently increased with increasing nominal concentrations and was stable over time. NaHCO_3_ addition resulted in slightly raised pH (from 7.9 to 8.1-8.2), which further increased over time (to 8.9-9 after 48 hours) but this was constant across the experimental salinity levels. Three replicate cohorts of 10 <24h old juveniles (2^nd^ to 4^th^ clutch) of animals grown under common-garden conditions were exposed in 210 ml jars containing 180 mL of a pre-set NaHCO_3_ and immobilisation was assessed after 48h as a proxy for mortality. Animals were considered dead when they did not show any muscular response after stirring the jar gently [48]. This design resulted in 21 populations x 6 clones x 8 salinity treatments x 3 replicates = 3024 experimental observations at 48h. To generate independent data for each replicate, the 126 lineages were cultured in such a manner that animals used for replicate observations were separated in time by at least two generations. When counting immobilised animals, we noticed that a limited number of cohorts contained 9 instead of 10 animals, and in two instances 8 and 7 individuals (56 cohorts of 3024 experimental units). Animals were not fed during the trial, in line with OECD guidelines of 48h toxicity testing assays [48]. Newborn *Daphnia* (<24h old) have been shown to perform limited to no grazing activity [49].

### Statistical analyses

#### Mortality upon salt exposures and evolution of salinity tolerance

All statistical tests were carried out using R version 4.3.1 [50]. We aimed to test for the impact of increased salinity levels on mortality and the signal of evolution in salinity tolerance among the clones isolated from bomb crater ponds and soda pans. Given the geographical distance and overall differences in salinity concentration between the two systems, we used a hierarchical approach, testing for the effect of region (bomb craters vs soda pans), populations within region, and clones within populations. We built hierarchical linear models for both raw data on mortality across all salinity treatments and for EC_50_ values (estimation methodology detailed in Supplementary Material, section Material and Methods).

Differences in salinity tolerance of *D. magna* cohorts across experimental salt concentrations (0-9 g L^−1^) were statistically tested using a generalised linear mixed-effect model (GLMM, package ‘lme4’, [51] using the number of dead and alive animals in each experimental unit as a binomial dependent variable, fitted via the insertion of a binomial error distribution with logit link function. Pond type (bomb crater vs soda pan), experimental salt concentration (continuous) and their interaction were included as explanatory variables. ‘Population’ (nested in ‘pond type’) and ‘clone’ (nested in ‘population’) were included as random effects. The model was tested and corrected for overdispersion (‘blmeco’ package, [52]; overdispersion function by [53]). The ‘car’ package [54] was used to estimate type II Wald χ^2^ statistics and p-values. Random effect ‘clone’ was investigated further via a model-comparison (full model, model without the random effect ‘clone’) to test for overall within-population variation in salt tolerance, indicative of overall evolutionary potential for salinity tolerance in populations. Marginal and conditional R^2^ values were calculated with the ‘MuMIn’ R package [55].

Half maximal effective concentration (EC_50_) offers an integrated way to assess tolerance to a chemical stressor by estimating a dose response function integrating data across all tested concentrations. We therefore also tested for adaptation to salinity among pond types and among populations within each of the two pond types applying linear mixed-effect models (package ‘lme4’), by regressing EC_50_ values against pond salinity (continuous, g L^−1^, at the time the clones were collected) and pond type. ‘Population’ was included as a random effect (nested in pond type). In order to have more reliable estimates of clone-specific EC_50_, we used all replicate data of a given clone (n = 3 per clone) to calculate one integrated value for EC_50_. As a result, no clonal random factor was added to the model as there was only one value per clone. A restricted maximum likelihood estimation (REML) method was used and degrees of freedom for fixed effects were corrected using the Kenward-Roger approximation [56]. P-values were computed using the ‘car’ package [54]. Model assumptions (normality, heterogeneity of variances across pond types) were visually checked (normality: plotting histograms of model residuals, residuals versus fitted values, and normal Q-Q plots) and formally tested (normality: Shapiro-Wilk test, heterogeneity of variances: Levene’s test with pond type as a grouping factor). No deviations with regard to model assumptions were found. Given the uneven spread along the pond salinity gradient of the two pond types, the same model was also run with log-transformed pond salinity, yielding qualitatively the same results (Supplementary Material, Table S2). In addition, linear mixed-effect regression models on EC_50_ values were run separately on the bomb crater pond and soda pan datasets, with pond salinity (continuous) as explanatory variable and ‘population’ included as a random effect, yielding qualitatively similar output (Supplementary Material, Table S3). We also tested for any potential underlying effect of spatial distances on differences in salinity tolerance using partial Mantel tests, applied separately for the two pond datasets (Supplementary Material, section Material and Methods).

#### Variation in salinity tolerance within populations

To explore whether variation in salinity tolerance within *D. magna* populations was a function of pond salinity and whether this reflects a weeding out of genotypes with lower salinity tolerance, we ran linear regression models. As response variables, we used the population level minimum and maximum EC_50_ values (i.e. the EC_50_ values of the clone of a given population with the lowest or highest salinity tolerance), the range in EC_50_ values (difference between population level minimum and maximum EC_50_ values), as well as the variance in EC_50_ values per population. All four models included pond salinity (continuous) as measured at the time of sampling the populations, pond type, and their interaction as explanatory variables. Model assumptions were visually checked and formally tested and were met in all models.

As we have multiple mortality data per clone (three replicate values per clone per exposure concentration), we can obtain a rough estimate of the amount of genetic variation for salt tolerance within populations and thus evolutionary potential using a clonal repeatability approach (quantifying the relative importance of among- and within-clone variation in salinity tolerance for each population). These are rough estimates because of the low number of replicate observations per clone and the low number of clones per population, but they provide a first indication of evolutionary potential. Given the observation that most variation in mortality was present between 5 and 7 g L^−1^ (below 5 g L^−1^ full survival, above 7 g L^−1^ quasi-full mortality, see Figure 2a), the amount of genetic variation in each population was calculated based on average clonal mortalities across salt exposure concentrations 5 to 7 g L^−1^ NaHCO_3_. Heritability as given by *h^2^ = V_g_ / V_g_+V_e_* was estimated for each of the 14 bomb crater pond and 7 soda pan populations separately following a clonal repeatability analysis and maximum-likelihood method [57,58]; we calculated among clonal variation (V_g_ + V_e_) and within clonal variation (V_g_) based on mean square model output values of among clonal variation (V_g_ + V_e_) and an error term (V_e_). We carried out a regression analysis on these rough estimates of within-population genetic variation calculated for each population with salinity observed in their habitat (continuous), pond type (bomb crater vs soda pan) and their interaction as explanatory variables. Model assumptions were visually checked and formally tested and were met in all models.

**Figure 2.**
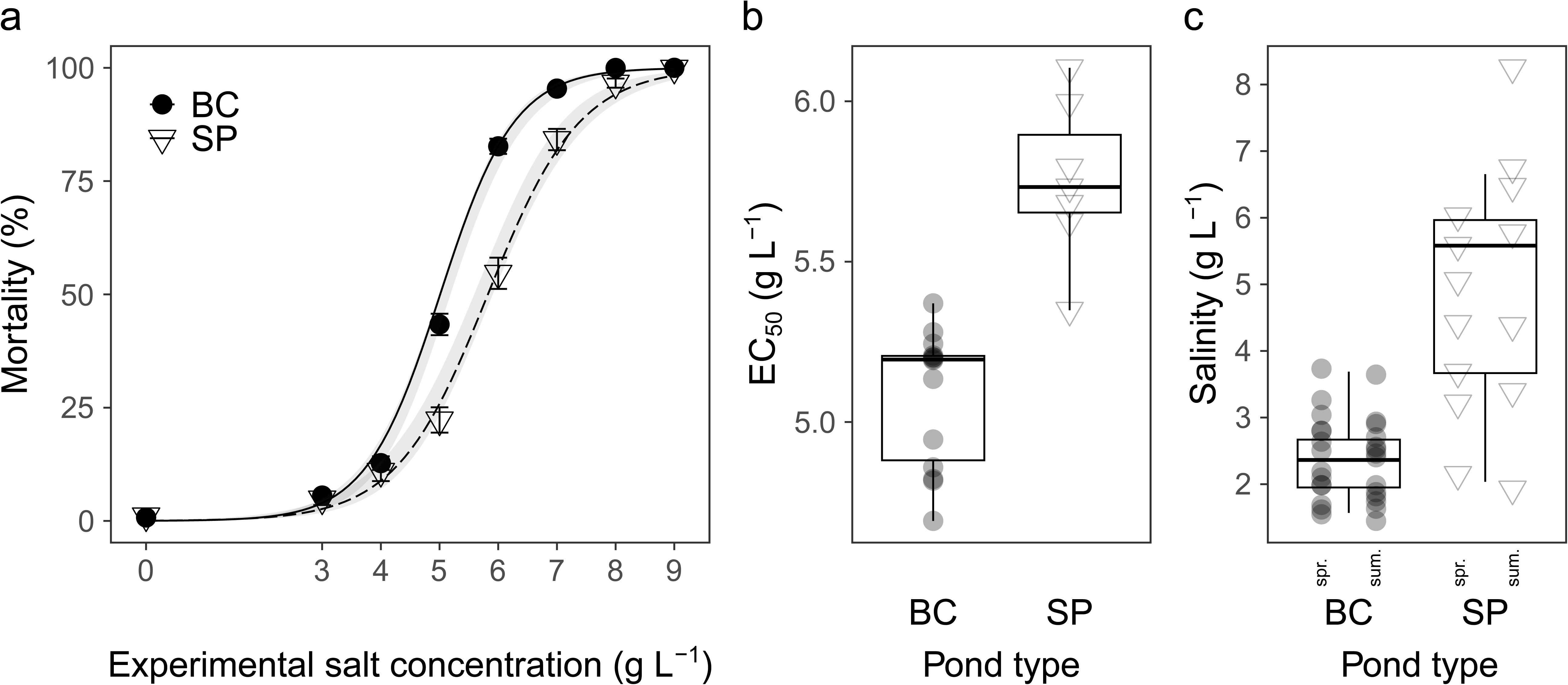
a) 48-h mortality (%, mean ± 1SE) of *Daphnia magna* from populations originating from bomb crater ponds (BC, N = 14) and soda pans (SP, N = 7), along the experimental salinity gradient. Trend lines represent fitted generalized linear mixed-effect models (with 95% confidence intervals). b) Salinity tolerance (EC_50_) of *D. magna* populations in the two pond types (boxplot with individual population mean EC_50_ values as overlayed symbols). c) Mean salinity of bomb crater ponds and soda pans (boxplots) which was averaged from mean spring (’spr.’) and summer (’sum.’) values (overlayed symbols) in each pond.

#### Nestedness pattern along the salinity gradient

There are in essence two ways in which populations can adapt to higher salinities: they can show a reduced occurrence or frequencies of genotypes with low salinity tolerance, or they can show a general shift in genotypes towards higher salinity tolerances. The former reflects a weeding out of sensitive genotypes and results in a ‘nested’ pattern where the range in sensitivity of genotypes in high salinity ponds is a subset of the range of sensitivity in low salinity ponds. The latter reflects a turnover type of process, resulting in an upward shift in sensitivities of genotypes without necessarily reducing the range in sensitivities. Given that we observed a strong correlation between minimum EC_50_ values and pond salinity in the bomb crater populations while no such correlation was observed for maximum EC_50_ values (see further), we tested for a nestedness pattern in salinity tolerance along the natural salinity gradient across habitats. Nestedness is typically tested in community data and uses occurrence data on taxa. In our analysis, we cannot use clone identities, because every individual likely belongs to a different clone. To develop a parallel analysis, we therefore binned the clones into functional identities (IDs) based on their EC_50_ values. We applied a manual binning to the full gradient of EC_50_ values in bomb craters, creating 6 classes of EC_50_ values with equal ranges (ID1: 3.73 - 4.11, ID2: 4.11 - 4.49, ID3: 4.49 - 4.87, ID4: 4.87 - 5.24, ID5:5.24 - 5.62, ID6: 5.62 - 6 g L^−1^). These categories were treated as functional IDs for clones exhibiting a gradient of EC_50_ across the ponds. We first tested whether their number in a local population decreases along the natural salinity gradient with a linear model. We then tested for the general presence of any nestedness pattern in the functional ID composition of the ponds with the functions ‘nestedchecker’ and ‘oecosimu’ in the ‘vegan’ package [59], using C-score statistics based on the null model method ‘r0’ with 2000 simulations. A nestedness plot was created using the ‘nestednodf’ function. Finally, we also tested whether the level of nestedness of individual pond populations (further referred as nestedness rank) corresponds to the natural salinity gradient with another linear model. For this model, we manually corrected the order of populations projected on the nestedness plot to include ties in the analysis, given that in our case, only presence-absence data was available, therefore the order of two ponds with the presence of the same functional identities is random on the nestedness plot. In contrast to the bomb crater ponds, the minimum EC_50_ values for soda pans did not show a significant relationship with pond salinity. Yet, to allow comparisons, we also carried out a parallel nestedness analysis for soda pans. If clone thinning is the mechanism consistent with increasing salinity, a positive relationship is expected between the nestedness rank of populations and salinity.

## RESULTS

### Mortality and evolution of salinity tolerance (EC_50_)

Mortality increased significantly with exposure to higher salinities, with a steep increase between 4 and 7 g L^−1^ (β ± SE: 2.820 ± 0.087 on log-odds scale, χ^2^ = 1346.712, p < 0.001; Figure 2a). The curve slopes were flatter for animals from soda pans (pond type x experimental salt concentration: χ^2^ = 20.238, p < 0.001; Figure 2a). This is paralleled by the overall higher salinity levels measured in the field for soda pans compared to bomb craters (Figure 1, Figure 2c). Adding a ‘clone’ random effect (nested within ‘population’) improved model fit (χ^2^ = 20.89, p < 0.001) indicating an overall effect of clonal variation within populations on survival.

Average EC_50_ values of *D. magna* were significantly higher for soda pan than for bomb crater pond populations (β ± SE: 1.166 ± 0.210, p < 0.001; Table 1, Figure 2b), consistent with their lower average mortalities observed at intermediate experimental salt concentrations (Figure 2a). Soda pan and bomb crater pond populations differed in how average EC_50_ values of populations varied as a function of pond salinity (pond type x salinity, β ± SE: −0.208 ± 0.075, p = 0.007; Table 1, Figure 3). Separate analysis on the two pond types confirmed that while average EC_50_ values of bomb crater pond populations increased significantly with pond salinity (β ± SE: 0.181 ± 0.060, p = 0.009; Figure 3, Supplementary Material, Table S3), no such signal of local adaptation was found in soda pans (β ± SE: −0.026 ± 0.056, p = 0.659; Figure 3, Supplementary Material, Table S3). Random variation attributed to population (nested in pond type) was not significant in any of the analyses (Table 1, Supplementary Information, Table S2, S3).

**Figure 3.**
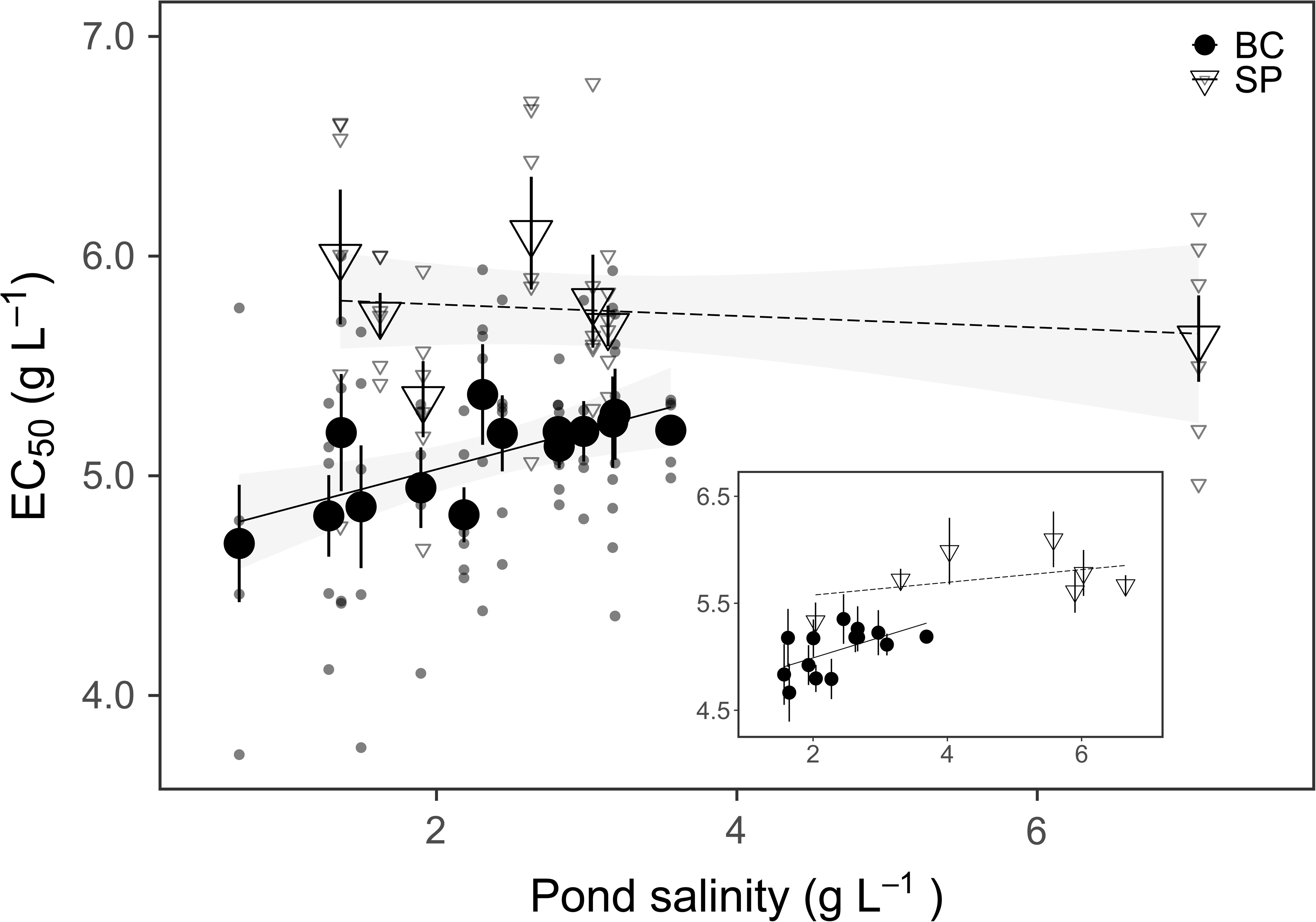
Salinity tolerance (mean EC_50_ ± 1SE, large symbols) of *Daphnia magna* populations originating from bomb crater ponds (BC, N = 14) and soda pans (SP, N = 7) as a function of pond salinity recorded in the field at the time the clones were collected. Trend lines illustrate linear models (with 95% confidence intervals). The small symbols show EC_50_ values of the six clones per population. The inset shows the relationship between salinity tolerance and mean salinity levels calculated from data from multiple years and seasons.

**Table 1.**
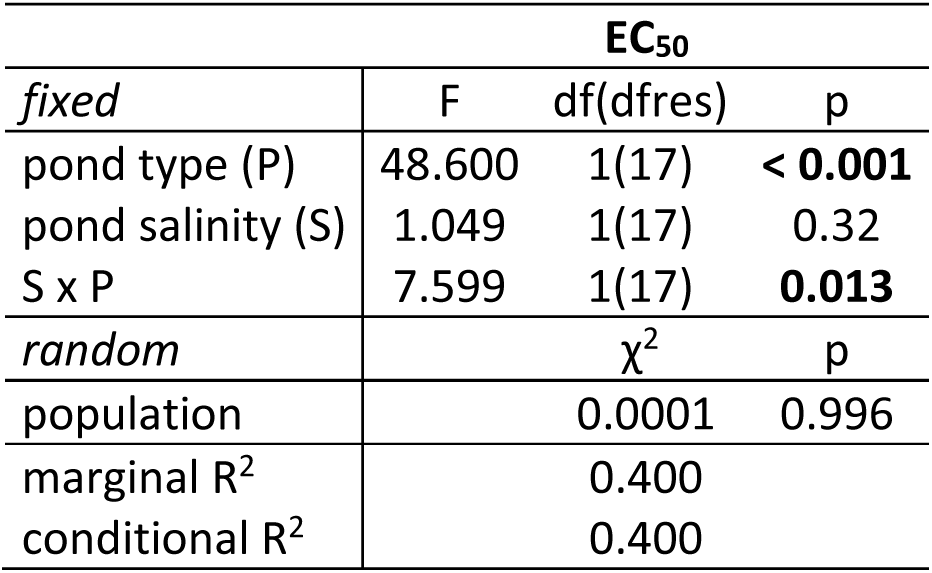
Anova model output of linear mixed-effect model on salinity tolerance (EC_50_) in response to pond salinity (g L^−1^), pond type (bomb crater pond vs soda pan) and their interaction, including a random effect for population (nested in pond type). Significant results (p < 0.05) are indicated in bold.

Pairwise differences in population mean EC_50_ values did not show a significant correlation with spatial distances either in the case of soda pans (partial Mantel’s r = 0.17, p = 0.201) or bomb crater ponds (r = 0.07, p = 0.291; Supplementary Material, Figure S1). We observed a significant relationship with pond salinity in bomb crater ponds (r = 0.46, p = 0.004), but not in soda pans (r = –0.23, p = 0.640).

### Variation in salinity tolerance within populations

Linear regression on the population-level range in clonal EC_50_ values revealed a significant effect of salinity x pond type (Supplementary Material, Table S4, Figure S2a), with a significant decrease with salinity only in the bomb crater ponds (β_BC_ ± SE: –0.420 ± 0.143, p = 0.009). The population-level variance in EC_50_ values was marginally significantly affected by salinity x pond type (Supplementary Material, Table S4, Figure S2), with a similar trend to that obtained for EC_50_ range (β_BC_ ± SE: –0.114 ± 0.044, p = 0.019). Within-population clonal maximum EC_50_ values were significantly higher in soda pans compared to bomb crater ponds (β_SP_ ± SE: = 0.694 ± 0.341, p = 0.058; pond type: F_1/17_ = 21.482, p < 0.001; Table 2, Figure 4a), and showed no relationship with pond salinity in either habitat type (Table 2, Figure 4a). The minimum EC_50_ values within each population showed a significant increase with increasing pond salinity in bomb crater ponds but not in soda pans (β_BC_ ± SE: 0.401 ± 0.089, p <0.001; Table 2, Figure 4b). Estimates of within-population genetic variation showed a trend for a negative association with salinity in bomb crater ponds but not in soda pans (β_SP_ ± SE: –0.183 ± 0.096, p = 0.073; Supplementary Material, Table S4, Figure S3).

**Figure 4.**
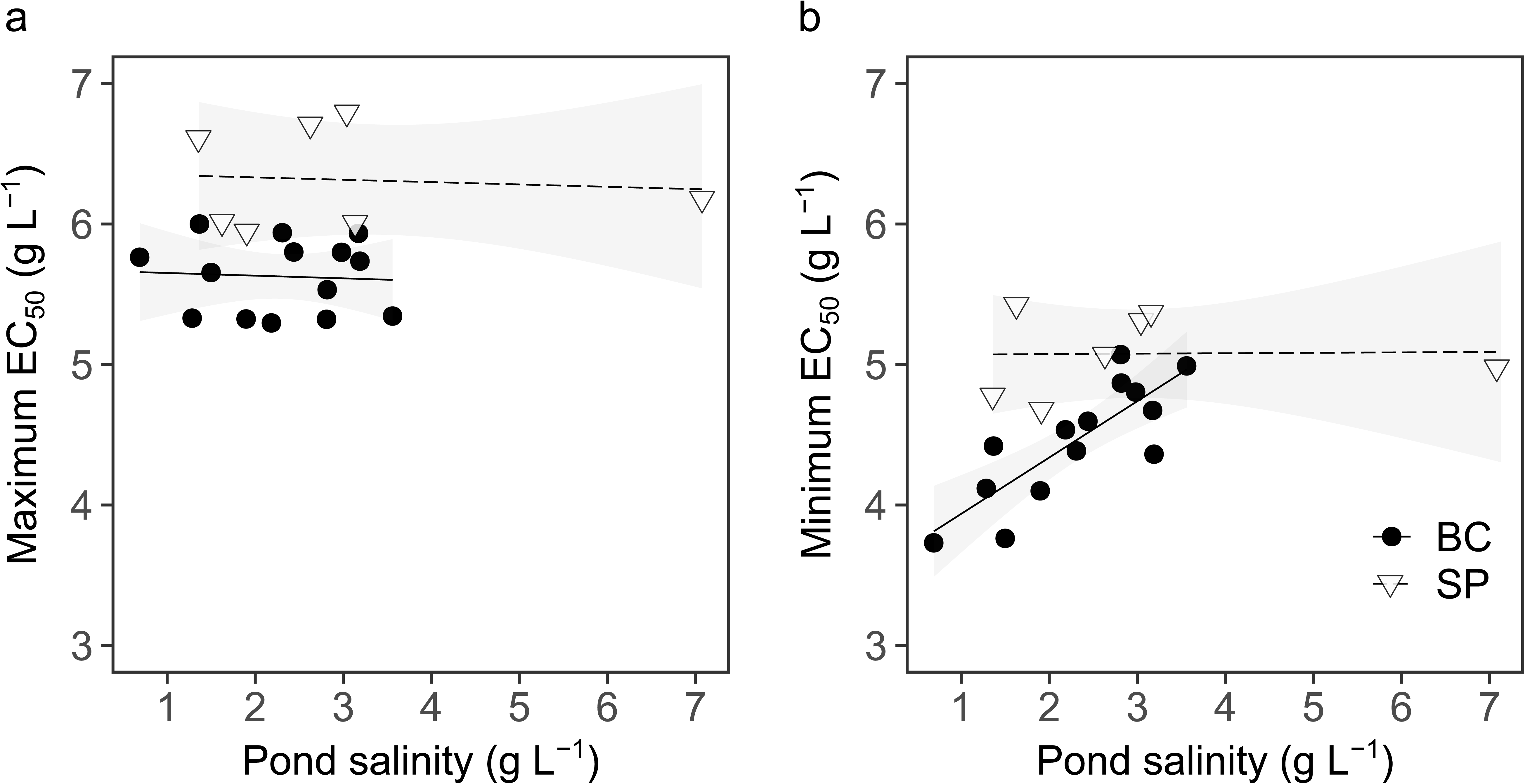
Maximum (a) and minimum EC_50_ (b) per population in bomb crater ponds (BC) and soda pans (SP), in response to pond salinity recorded in the field at the time the clones were collected (linear models with 95% confidence intervals).

**Table 2.** Anova model output of linear models on within-population clonal level minimum and maximum EC_50_ in response to pond type (bomb crater pond vs soda pan) and pond salinity (g L^−1^). Significant results (p < 0.05) are indicated in bold.

|  | minimum EC<sub>50</sub> |  |  | maximum EC<sub>50</sub> |  |  |
| --- | --- | --- | --- | --- | --- | --- |
|  | F | df(dfres) | p | F | df(dfres) | p |
| pond type (P) | 16.699 | 1(17) | **0.007** | 21.482 | 1(17) | **<0.001** |
| pond salinity (S) | 6.168 | 1(17) | **0.024** | 0.097 | 1(17) | 0.759 |
| S x P | 13.906 | 1(17) | **0.002** | <0.001 | 1(17) | 0.984 |
| adjusted R<sup>2</sup> | 0.670 |  |  | 0.490 |  |  |

### Nestedness pattern along the salinity gradient

Local diversity in salinity tolerance (measured as the number of functional IDs per population) showed a marginally significant negative trend with pond salinity in the bomb craters (R^2^ = 0.229, p = 0.083; Figure 5a). Here, a significant nestedness pattern was also detected (C-score = 7.4, p < 0.01; Figure 5b), and nestedness rank showed a marginally significant trend corresponding to the salinity gradient (R^2^ = 0.225, p = 0.088; Figure 5c). In soda pans, none of these patterns were significant (local no. of functional IDs vs pond salinity: R^2^ = 0.359, p = 0.155, nestedness: C-score = 1.7, p = 0.102, nestedness rank vs pond salinity: R^2^ = 0.400, p = 0.128; Supplementary Material, Figure S3).

**Figure 5.**
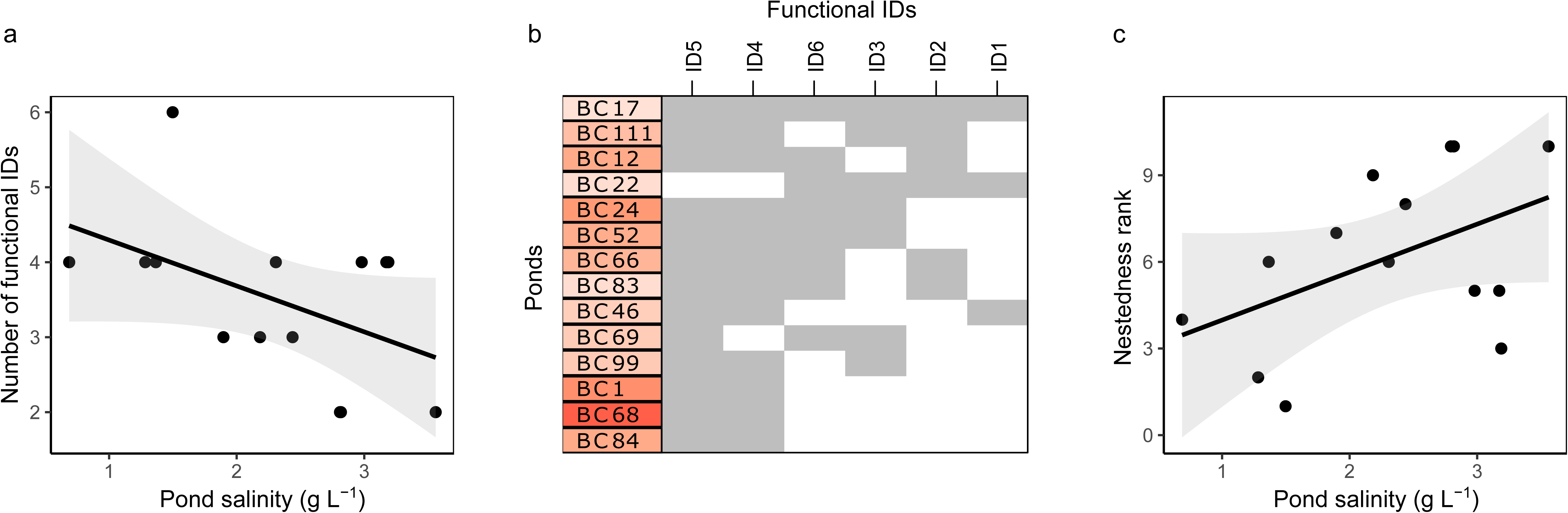
a) The observed number of functional identities (IDs; genotypes belonging to six classes of EC_50_ salinity tolerance values with equal ranges) along pond salinity in the 14 bomb crater ponds at the time the clones were collected (linear model with 95% confidence intervals). b) Nestedness plot based on the six functional IDs (ID1 - ID6, ranging from high salinity sensitivity to high salinity tolerance) in the 14 bomb crater populations, with grey shading showing the presence of functional IDs in each population; ponds are given colour codes visualising mean pond salinities (using the same colour gradient as in Figure 1). c) Relationship between the nestedness rank of populations (lowest rank being the population with the most complete representation of functional IDs) and pond salinity in the bomb crater ponds at the time the clones were collected (linear model with 95% confidence intervals).

## DISCUSSION

Our results clearly show a strong and significant pattern of both regional genetic differentiation between the two study systems, as well as local genetic differentiation in salinity tolerance only in the bomb crater ponds. Both of these patterns are in line with predictions of adaptive evolution. First, the higher average salinity tolerance of soda pan populations is associated with their generally higher regional salinity levels compared to the bomb crater ponds. Second, the salinity tolerance of bomb crater pond populations is positively correlated with the salinities of their local habitats. This pattern represents a striking case of local adaptation at very small spatial scales, as these ponds are located only a few to a few hundred metres from each other. These results also show that the degree to which local adaptation is observed is not necessarily associated with geographic distance [7], as the soda pans are geographically much more distant to each other than the bomb crater ponds. Finally, we also show that the range in clonal values for salinity tolerance in bomb crater ponds is higher in populations with low salinities. This is driven by the minimum clonal value of salinity tolerance within a population significantly increasing with pond salinity.

Patterns of local adaptation have been reported for a wide range of ecosystems and at wide spatial scales [7,60]. Earlier studies have provided ample evidence for *Daphnia* to have the capacity to genetically adapt to a wide range of abiotic and biotic stressors (e.g. fish predation [61,62]; algal toxins [63,64]; parasites [65]; warming [66,67]; urbanisation [68,69]; pollutants [70,71]). Earlier studies, mostly focusing on contrasting two populations inhabiting ponds or lakes with different salinities, have also shown that salinity tolerance levels of *Daphnia* clones and populations reflect the salinity of their habitats [72,73]. Studies on obligately parthenogenetic *D. pulex* showed clonal sorting along salinity gradients in ponds in the Churchill region of northeastern Manitoba, Canada [42,43]. Experimental evolution revealed local adaptation of *Daphnia* to a moderate increase in salinity within 2.5 months, representing an estimated 5-10 generations of parthenogenetic reproduction. This reflects fast clonal sorting during the parthenogenetic phase, highlighting the potential for adaptation within one growing season [40]. Our study quantified local genetic differentiation in salinity tolerance of 21 *D. magna* populations in relation to habitat salinity in natural landscapes of both closely-spaced bomb crater ponds and geographically more distant soda pans. Given the extent and pace of freshwater salinisation, these results are important as they indicate that *Daphnia* populations may adapt at different spatial scales including ponds being separated by just a few tens of metres (e.g., one population pair that showed one of the highest differences in EC_50_ were <100 m distance from each other, see Figure S1). Given their important role as grazers and prey in the sodic waters we studied [34,37,74,75], it is likely that this level of local adaptation will have important ecosystem implications [76].

In line with our initial expectation, the pattern of regional genetic differentiation between the soda pans and the bomb crater ponds reflects the differences in salinity ranges between the two types of habitats. Soda pans are very shallow and relatively large in surface area compared to the bomb crater ponds, resulting in much higher surface to volume ratios. As a result, most soda pans dry up yearly [31,77], unlike a large share of bomb crater ponds that only dry up in exceptionally dry years (C.F. Vad, pers. obs.). Consequently, overall salinity levels in the soda pans are higher than in the bomb crater ponds and especially show higher peaks (Figure 2, Supplementary Material, Table S1). Our results indicate that the *Daphnia* populations studied have regionally adapted to these habitat-specific differences in salinity dynamics.

We observe a pattern of local adaptation in the bomb craters but not in the soda pans. While local genetic differentiation in the bomb crater ponds occurs at a very small spatial scale to at most a few hundreds of metres, we do not observe local adaptation among soda pans, even though distances between them range up to almost 20 km. There are multiple potential explanations for this counter-intuitive difference. First, soda pans show a higher amount of within- and among-year differences in salinity levels, and thus a higher ratio of within compared to among habitat variation in salinity levels than in the bomb craters. This reduces the degree of differential selection while increasing the overall strength of selection for high salinity tolerances. Second, while geographic distances among soda pans are larger, it is not certain that this translates into more gene flow among bomb craters than among soda pans. Their large surface areas, the fact that they have a dry phase exposing dormant eggs to wind every year, and the fact that they attract large amounts of waterbirds [19,45] and are exposed to strong directional winds [44] all promote dispersal of *Daphnia* among soda pans. In contrast, the bomb crater ponds are sheltered from wind due to their lower surface-to-volume ratio further enhanced by reedbelt in many of them and are too small to attract large numbers of waterbirds. Furthermore, they are also physically more isolated due to their elevated rim created during the explosion of the original bombs (Supplementary Material, Figure S5). This is expected to reduce their connectivity through dispersal even under close geographic distances, which is also a prevalent pattern in the structure of their zooplankton metacommunities [78].

In addition to the observed EC_50_ population means that strongly indicate local adaptation in the bomb crater system, our data show substantial overall inter-clonal variation in mortality upon experimental salt exposure within populations of both bomb craters and soda pans. In the case of the bomb crater ponds, the additional analyses on salinity tolerance ranges, minima and maxima reveal a lack of clones with low salinity tolerance in the more saline bomb crater ponds, while clones with maximum salinity tolerance show no relationship with salinity. Consequently, there is a significant reduction of the population-level salinity tolerance range with increasing pond salinity. This implies that selection against clones with low salinity tolerances in high salinity ponds is stronger than selection against clones with high salinity tolerances in low salinity ponds. The presence of clones with high salinity tolerance in less saline ponds might reflect their occasional success in extreme years. The broad range in salinity tolerances we observe in low salinity ponds has important consequences, as it is expected to facilitate rapid adaptation to higher salinity stress also in the low salinity ponds. These findings were further supported by our observation that estimates of within-population genetic variation (based on clonal repeatability analyses of mortality between 5-7 g L^−1^ of experimental salt concentrations) tended to decline with increasing pond salinity in the bomb crater pond but not in the soda pan populations.

The nestedness analysis we performed confirmed the pattern suggested by the significant relationship between minimum salinity tolerance and pond salinity. Our analysis indeed shows a striking nested pattern, with a tendency of the presence of a more diverse set of salinity tolerances at lower salinities, with tolerance levels thinning to clones of higher tolerances (identified by higher IDs on the nestedness plot; Figure 5b) in more saline bomb crater ponds. This result parallels the observation in community ecology that nestedness is a general pattern along salinity gradients [14,23,24]. Such types of analyses, originated in community ecology, bring valuable insights into the mechanisms of local adaptation. The pattern observed for within-population genetic variation and for the association between nestedness rank and observed salinities in the habitats are only indicative trends as they are marginally significant. But they are further supporting the significant association between minimum salinity tolerance of clones in populations and observed salinities in the field.

We studied seven soda pan and fourteen bomb crater pond populations. The absence of evidence for local adaptation in the soda pans is, however, not due to a lower power of our analyses for these systems. Indeed, Figure 3 shows no trend for a positive association between salt tolerance and salinity measured in the field for the soda pans, while such an association would be readily apparent even with less bomb crater ponds. Pond salinities used in the analyses were quantified at the same moment the clones were sampled from the field, during an early season snapshot sampling in a relatively wet year, resulting in relatively low salinities. Salinities in these systems vary across seasons and years (see Figure 2c). Yet, the Figure 3 inset shows a similar pattern when we plot salinity tolerances against the average value of salinities measured during several field campaigns spread across years. Finally, typical for field-based studies, salinity may also co-vary with other environmental dimensions in these habitats, e.g., with pH [14,34]. While some influence of confounding factors cannot be excluded, the positive association between population-specific tolerances to a specific stressor, salinity, as observed in the laboratory (with the absence of pH gradient across experimental treatments) and differences in that same stressor in the studied habitats provides a strong indication for differences in salinity being the driving force of genetic differentiation. It is well-known that salinity is an important stressor that in a very direct way affects physiology and cellular processes [12]. It is less straightforward to explain how a change in salinity tolerance would be driven by a gradient in pH or other environmental gradients. The observation that mortality in the laboratory increased strongly between salinity levels of 5 and 8 g L^−1^ supports our interpretation, as we know from earlier studies that *D. magna* only occurs in the studied systems up to salinity levels of approximately 9 g L^−1^ [14].

Salinity is a major abiotic stress factor for freshwater biota strongly impacting community composition and diversity [13,24,25]. Salinisation is an important component of global change with several drivers increasing salinity in inland waters [20,21]. Our results show that *D. magna* can locally adapt to differences in natural salinity level which may imply a potential for adaptation to anthropogenic salinisation. Our results extend earlier work on evolution of salinity tolerance in *Daphnia* by quantifying adaptation to natural salinity gradients in two sets of contrasting habitats, showing how regional and local adaptation integrate to shape patterns of salinity tolerance. We also show that local adaptation is achieved mainly by a weeding out of clones with low salinity tolerances rather than by a shift in both minimum and maximum clonal tolerances within populations. This nested pattern results in a broad range of salinity tolerances of clones in low salinity habitats, while only a high salinity tolerance subset is present in the high salinity populations. These results suggest that most populations harbour genetic variation in salinity tolerance, indicating evolutionary potential for quick adaptation to salinity changes, and that increased salinity in bomb crater ponds results in local adaptation, with low tolerant genotypes being weeded out, resulting in a tendency for a reduction in the amount of within-population genetic variation.

Given the important ecological roles of large-bodied *Daphnia* [39,79] and their dominance in the study systems [14,34,37], it is our prediction that regional and local adaptation may have important implications for the ecology of the study systems. Quantifying these effects would require integrated field transplant experiments using animals grown under common garden conditions. Our results suggest that this impact of local adaptation on ecosystem processes will likely be especially strong in the bomb crater ponds, despite their close geographic proximity.

## Acknowledgements

The study was supported by the FWO project no. G0C3818N and KU Leuven Research Council project C16/2017/002. CFV and ZH acknowledge further support by the NKFIH project no. 2019-2.1.11-TÉT-2020-00159. ZH was furthermore supported by the János Bolyai Research Scholarship of the Hungarian Academy of Sciences (Grant number BO/00392/20/8). KIB acknowledges further support by KU Leuven PDM/18/112 and FWO project no. 1222120N. The authors thank Edwin van den Berg, Ria Van Houdt, Geert Neyens, Rony Van Aerschot, and Christian Preiler for their help during the extensive laboratory experiments.

