## Supplementary Material for "Regional and fine-scale local adaptation in salinity tolerance in *Daphnia* inhabiting contrasting clusters of inland saline waters"

**Material and Methods**

*Culturing* Daphnia *for purging maternal effects*

A parthenogenetically produced offspring of each clone was used to inoculate 6 independent monoclonal cultures (n = 126 x 6) consisting of one *D. magna* individual in a 210-mL jar containing 180 mL of dechlorinated tap water and kept under standardised laboratory conditions (climatized walk-in room 20°C ± 1°C; 14:10h Light:Dark photoperiod, 50% medium renewal three times a week) and fed daily by the unicellular batch-cultured green algae *Acutodesmus obliquus* ($10^5$ cells $mL^{-1}$, ~ 1 mg C $L^{-1}$). All lineages were cultured for at least two generations under these standardised conditions to reduce interference from (grand)maternal effects. At the start of each new generation, the culture was re-inoculated with two randomly picked second clutch offspring juveniles reared until sub-adult stage, after which rearing continued with one individual. At the start of the third generation, 15 newborn second clutch juveniles (<24h old) were inoculated in 500-mL jars filled with dechlorinated tap water to start up cultures that would generate experimental neonates under standardised conditions. These cultures were transferred to a 20°C temperature-controlled water bath and kept under the same culture conditions and feeding regime as described above. Their densities were retained at 15 individuals per 500 ml jar, and they represent the mother generation from which experimental animals were generated.

*Estimating $EC_{50}$*

Based on the three clonal replicates of *D. magna* mortality counts (after 48h) from each laboratory salt concentration, the $EC_{50}$ (half maximal effective concentration; $NaHCO_3$ concentration at which 50% of the animals die, [1]) was determined for each bomb crater and soda pan genotype. A general linear model with binomial error distribution and logit link function was fitted to predict mortality percentages of each clone for experimental salt concentrations which ranged from 0 to 9 with an interval of 0.0001.

*Relating among-population differences in tolerance to spatial distance and to differences in salinity*

To test whether genetic differentiation is a function of geographic distances, we tested the relationship between pairwise spatial distances among habitats and pairwise differences in mean $EC_{50}$ of populations. This was done via partial Mantel tests (999 permutations) by first partialling out the effect of salinity in the models. We then also tested genetic differentiation as a function of differences in field salinities via a second set of partial Mantel tests when spatial distances were partialled out. Partial Mantel tests were performed and Mantel correlation coefficients were calculated with the function 'partial.mantel' in 'vegan' [2], using Euclidean distances for mean $EC_{50}$ and field salinity, while spatial distances were calculated with 'rdist.earth' in the 'fields' package [3].

*References*


1. OECD. 2004 Test No. 202: *Daphnia* sp. Acute Immobilisation Test. In *OECD Guidelines for the Testing of Chemicals, Section 2*, pp. 1-12. Paris: OECD Publishing.
2. Oksanen J *et al.* 2022 vegan: Community Ecology Package.
3. Nychka D, Furrer R, Paige J, Sain S. 2023 fields: Tools for Spatial Data.

Table S1 – Geographical coordinates and environmental characteristics of bomb crater ponds and soda pans from which the experimental populations of *Daphnia magna* originated. Environmental variables measured in spring 2018 during the collection of *Daphnia magna* populations are presented along with surface area and mean spring and summer salinity values. For more details on specific sample analysis procedures and protocols, we refer to [4] and [5]. Mean spring and summer salinities were calculated as averaged salinity values of multiple years (spring data: April 2014, 2015, and 2018 for bomb crater ponds, while April 2010, 2014 and 2018 for soda pans; summer data: mid-June 2014 and 2015 for bomb crater ponds, mid-June 2009 for soda pans). Mean salinity was then calculated as the mean of these two values. IDs of bomb crater ponds are numbers given by the authors at the first mapping of all ponds in 2014. Abbreviations for soda pans: APME - Apetloner Meierhoflacke, HERR - Herrnsee, MSTI - Mittlerer Stinkersee, OWOR - Östliche Wörthenlacke, RUND - Runde Lacke, SECH - Sechsmadlacke, ZICK - Zicklacke.

| | | | | **Environmental characteristics in 2018** | | | | | | | | |
|---|---|---|---|---|---|---|---|---|---|---|---|---|
| **Habitat type** | **ID** | **latitude** | **longitude** | **salinity ($g L^{-1}$)** | **TSS ($mg L^{-1}$)** | **chlorophyll *a* ($\mu g L^{-1}$)** | **pH** | **water depth (cm)** | **surface area ($m^2$)** | **mean spring salinity ($g L^{-1}$)** | **mean summer salinity ($g L^{-1}$)** | **mean salinity ($g L^{-1}$)** |
| Bomb crater ponds | BC1 | 47° 7'16.03" | 19° 8'0.37" | 2.82 | 12.0 | 0.8 | 9.1 | 50 | 78.5 | 3.3 | 2.9 | 3.1 |
| | BC12 | 47° 7'26.60" | 19° 8'2.09" | 3.14 | 12.4 | 8.4 | 8.9 | 65 | 50.2 | 2.8 | 2.5 | 2.7 |
| | BC17 | 47° 7'26.75" | 19° 8'6.44" | 1.50 | 6.9 | 1.6 | 8.5 | 78 | 33.2 | 1.7 | 1.4 | 1.6 |
| | BC22 | 47° 7'35.64" | 19° 8'7.30" | 0.69 | 20.8 | 1.0 | 8.3 | 70 | 50.2 | 1.5 | 1.7 | 1.6 |
| | BC24 | 47° 7'34.70" | 19° 8'10.86" | 3.17 | 21.2 | 0.6 | 8.9 | 70 | 50.2 | 3.0 | 2.9 | 3.0 |
| | BC46 | 47° 7'21.56" | 19° 8'12.69" | 1.90 | 4.4 | 2.1 | 8.5 | 70 | 63.6 | 2.0 | 1.9 | 1.9 |
| | BC52 | 47° 7'24.54" | 19° 8'8.72" | 2.98 | 7.0 | 0.3 | 8.9 | 68 | 28.3 | 2.8 | 2.5 | 2.6 |
| | BC66 | 47° 7'27.80" | 19° 8'16.77" | 2.31 | 6.5 | 0.3 | 9.0 | 60 | 78.5 | 2.5 | 2.4 | 2.5 |
| | BC68 | 47° 7'27.34" | 19° 8'15.97" | 3.56 | 19.2 | 0.4 | 9.0 | 68 | 113.0 | 3.7 | 3.6 | 3.7 |
| | BC69 | 47° 7'26.89" | 19° 8'15.38" | 2.44 | 7.6 | 1.6 | 8.7 | 76 | 38.5 | 2.2 | 1.8 | 2.0 |
| | BC83 | 47° 7'21.28" | 19° 8'23.03" | 1.37 | 2.5 | 0.7 | 8.7 | 75 | 63.6 | 1.6 | 1.6 | 1.6 |
| | BC84 | 47° 7'20.93" | 19° 8'22.32" | 2.81 | 7.8 | 0.6 | 8.9 | 70 | 86.5 | 2.6 | 2.7 | 2.7 |
| | BC99 | 47° 7'18.26" | 19° 8'18.00" | 2.18 | 5.2 | 0.5 | 8.6 | 80 | 70.8 | 2.1 | 2.0 | 2.0 |
| | BC111 | 47° 7'13.19" | 19° 8'29.75" | 1.28 | 3.6 | 0.8 | 8.7 | 65 | 63.6 | 2.0 | 2.6 | 2.3 |
| Soda pans | APME | 47°43'17.67" | 16°49'26.55" | 3.14 | 125.7 | 37.9 | 9.1 | 35 | 4450 | 5.1 | 8.3 | 6.7 |
| | HERR | 47°44'40.43" | 16°46'11.96" | 7.07 | 64.9 | 43.8 | 9.2 | 31 | 1350 | 6.0 | 5.8 | 5.9 |
| | MSTI | 47°48'24.33" | 16°47'15.08" | 2.63 | 1030.0 | 131.3 | 9.2 | 24 | 4400 | 4.4 | 6.7 | 5.6 |
| | OWOR | 47°46'24.04" | 16°52'48.34" | 1.91 | 22.2 | 2.7 | 8.8 | 44 | 23300 | 2.1 | 1.9 | 2.0 |
| | RUND | 47°47'8.68" | 16°47'33.88" | 3.04 | 524.0 | 82.3 | 9.1 | 30 | 3260 | 5.6 | 6.5 | 6.0 |
| | SECH | 47°47'1.64" | 16°53'2.80" | 1.36 | 574.0 | 56.2 | 8.9 | 21 | 9300 | 3.7 | 4.4 | 4.0 |

| | | | | | | | | | | | |
|---|---|---|---|---|---|---|---|---|---|---|---|
| ZICK | 47°46’1.07” | 16°47’5.11” | 1.63 | 265.3 | 33.1 | 9.2 | 26 | 190380 | 3.2 | 3.4 | 3.3 |

*References*


4. Horváth Z, Vad CF, Tóth A, Zsuga K, Boros E, Vörös L, Ptacnik R. 2014 Opposing patterns of zooplankton diversity and functioning along a natural stress gradient: when the going gets tough, the tough get going. *Oikos* **123**, 461–471. (doi:10.1111/j.1600-0706.2013.00575.x)

5. Vad CF *et al.* 2017 Wartime scars or reservoirs of biodiversity? The value of bomb crater ponds in aquatic conservation. *Biol. Conserv.* **209**, 253–262. (doi:10.1016/j.biocon.2017.02.025)

**Results**

**Figures**

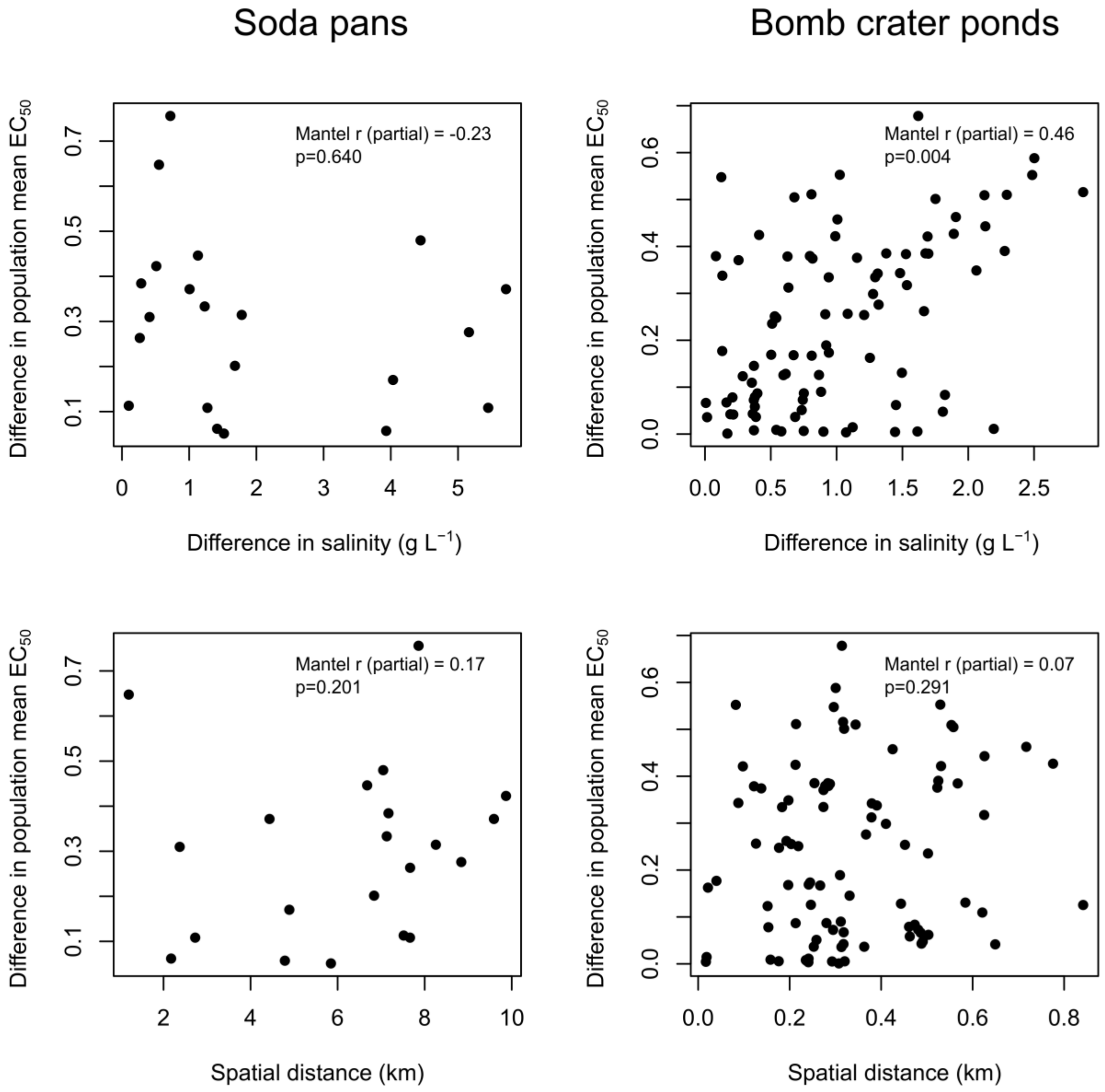


Fig S1 – The relationship between among-population differences in salinity tolerance (based on differences in mean $EC_{50}$) and pairwise differences in salinity (at the time the clones were collected; upper row) and pairwise spatial distances among habitats, shown separately for soda pans (left) and bomb crater ponds (right).

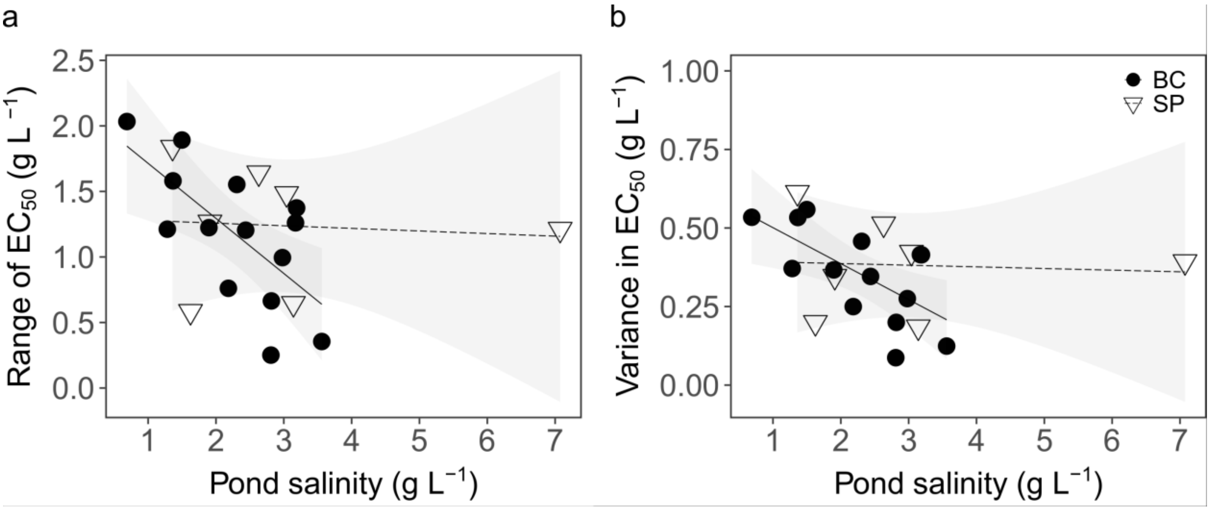


Fig. S2 – a) Observed population-level range in $EC_{50}$ values, calculated as the difference between the clone with the highest tolerance of the given population and the one with the lowest tolerance in that population, and b) variance in population-level $EC_{50}$ values per population in bomb crater ponds (BC) and soda pans (SP), in response to pond salinity recorded in the field at the time the clones were collected (linear models with 95% confidence intervals).

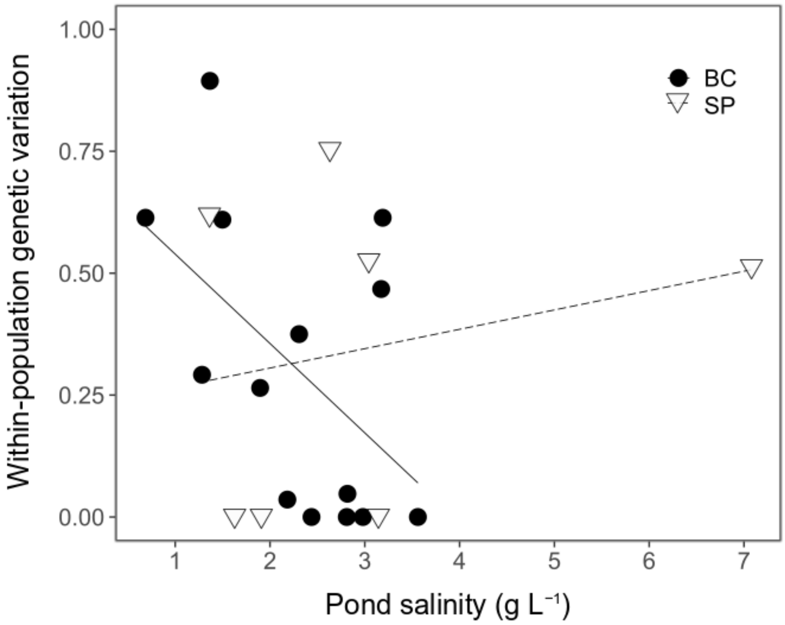


Fig. S3 – Within-population genetic variation for all studied bomb crater ponds (BC) and soda pans (SP) against pond salinity recorded in the field at the time the clones were collected (trend lines are from linear models).

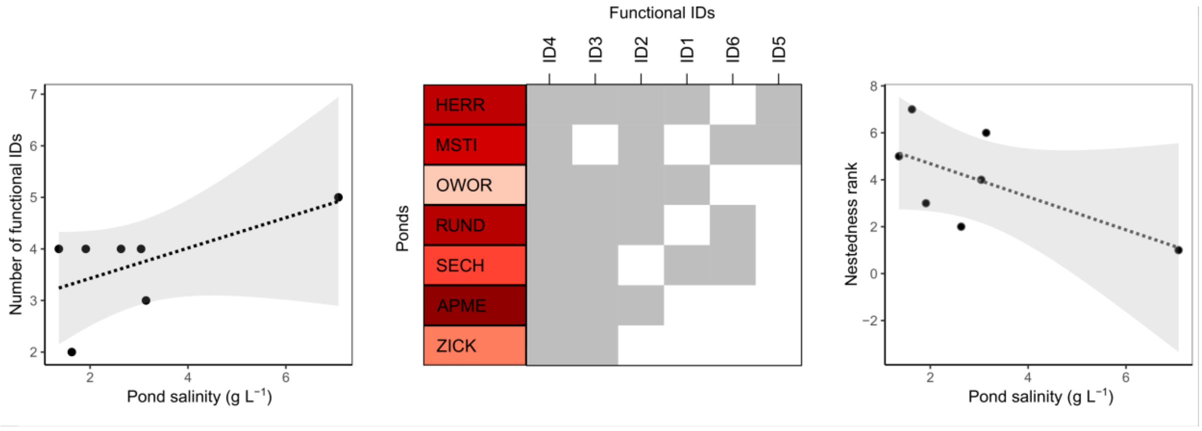


Fig. S4 – a) The observed number of functional identities (IDs; genotypes belonging to six classes of $EC_{50}$ salinity tolerance values with equal ranges) along pond salinity in the 7 soda pans at the time the clones were collected (linear model with 95% confidence intervals). b) Nestedness plot based on the six functional IDs (ID1 - ID6, ranging from high salinity sensitivity to high salinity tolerance) in the 7 soda pan populations (see Table S1 for names), with grey shading showing the presence of functional IDs in each population. Soda pans are given colour codes visualising mean pond salinities (using the same colour gradient as in Figure 1). c) Relationship between the nestedness rank of populations (lowest rank being the population with the most complete representation of functional IDs) and pond salinity in the soda pans at the time the clones were collected (linear model with 95% confidence intervals).

**Tables**

Table S2 – Anova model output of a linear mixed-effect model on salt tolerance ($EC_{50}$) in response to pond salinity (log transformed, g $L^{-1}$), pond type (bomb crater vs. soda pan), and their interaction, including a random effect for population (nested in pond type). Significant results ($p < 0.05$) are indicated in bold, while marginally significant ones (<0.1) in italics.

| | | **$EC_{50}$** | |
|---|---|---|---|
| ***fixed*** | **F** | **df(dfres)** | **p** |
| pond type (P) | 48.287 | 1(17) | **< 0.001** |
| log(salinity (S)) | 3.667 | 1(17) | *0.072* |
| log (S) x P | 5.349 | 1(17) | **0.034** |
| ***random*** | | **χ2** | **p** |
| population | | <0.001 | 1.000 |
| marginal $R^2$ | | 0.341 | |
| conditional $R^2$ | | 0.341 | |

Table S3 – Anova model output of linear mixed-effect model on salt tolerance ( $EC_{50}$) in response to pond salinity (g $L^{-1}$) for both a) bomb crater pond and b) soda pan datasets separately, including a random effect for population. Significant results ($p < 0.05$) are indicated in bold.

| | | **$EC_{50}$** | |
|---|---|---|---|
| | **F** | **df(dfres)** | **p** |
| **a) Bomb craters** | | | |
| pond salinity | 10.116 | 1(12) | **0.011** |
| *population* | | | 1.000 |
| marginal $R^2$ | | 0.099 | |
| conditional $R^2$ | | 0.099 | |
| **b) Soda pans** | | | |
| pond salinity | 0.2193 | 1(5) | 0.659 |
| *population* | | | 0.401 |
| marginal $R^2$ | | 0.008 | |
| conditional $R^2$ | | 0.112 | |

Table S4 – Anova model output of linear models on within-population level range and variance in clonal $EC_{50}$ values as well as within-population genetic variation in response to pond type (bomb crater vs. soda pan) and pond salinity (g $L^{-1}$). Significant results ($p < 0.05$) are indicated in bold while marginally significant ones ($p < 0.1$) in italics.

| | **$EC_{50}$ range** | | | **$EC_{50}$ variance** | | | **Within-population genetic variation** | | |
|---|---|---|---|---|---|---|---|---|---|
| | **F** | **df(dfres)** | **p** | **F** | **df(dfres)** | **p** | **F** | **df(dfres)** | **p** |
| pond type (P) | 0.119 | 1(17) | 0.735 | 0.713 | 1 (17) | 0.410 | 0.099 | 1(17) | 0.757 |
| pond salinity (S) | 3.173 | 1(17) | *0.093* | 2.473 | 1 (17) | 0.134 | 0.255 | 1(17) | 0.620 |
| S x P | 5.534 | 1(17) | **0.031** | 4.346 | 1 (17) | *0.053* | 3.813 | 1(17) | *0.067* |
| adjusted $R^2$ | 0.270 | | | 0.168 | | | 0.055 | | |

**Discussion**

Fig. S5 – Bomb crater ponds during sampling in April 2018 (left) and in an extremely wet season in March 2010 (right). The elevated crater rims visible around their perimeters are results of the original explosions, sheltering the ponds from strong winds and any potential hydrological connectivity. Photo credits: Luc De Meester (left), Zsófia Horváth (right).

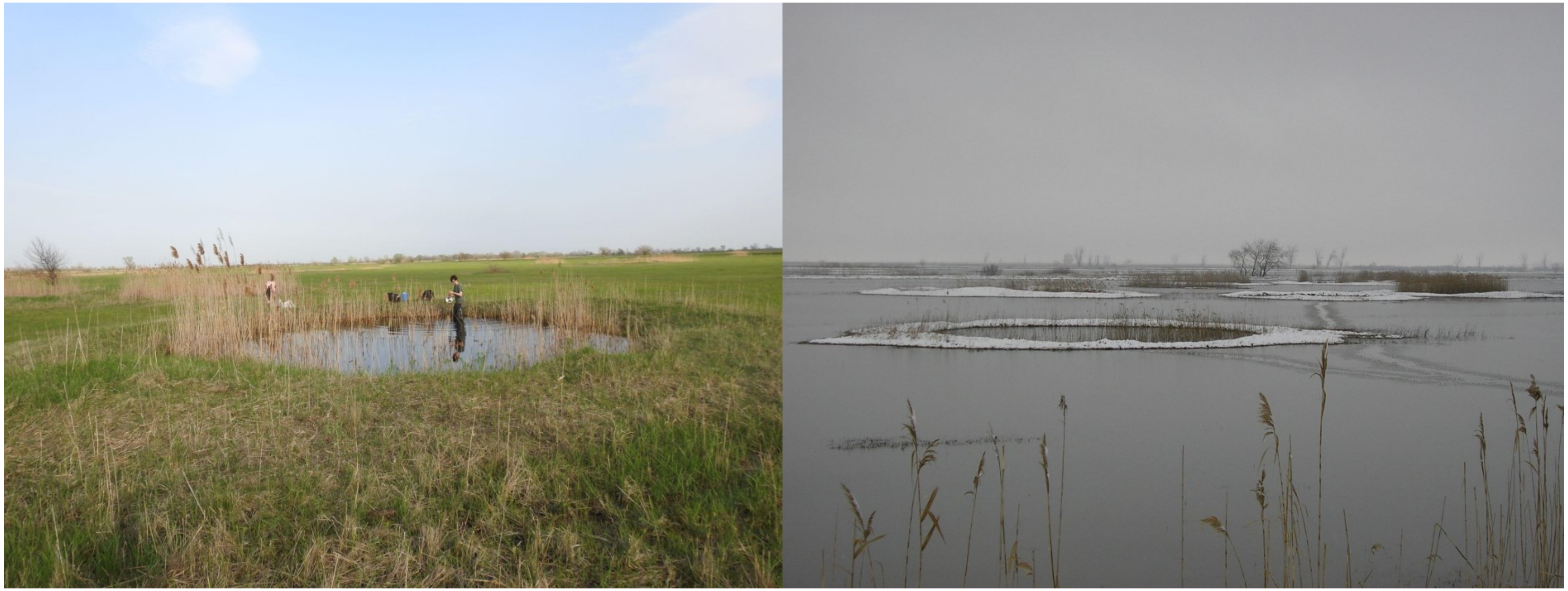
